# The genome assembly of *Rhabditoides inermis* from a complex microbial community reveals further evidence for parallel gene family expansions across multiple nematodes

**DOI:** 10.1101/2024.08.02.605984

**Authors:** Christian Rödelsperger, Waltraud Röseler, Marina Athanasouli, Sara Wighard, Matthias Herrmann, Ralf J. Sommer

## Abstract

**Background:** Free-living nematodes such as *Caenorhabditis elegans* and *Pristionchus pacificus* are powerful model systems for linking specific traits to their underlying genetic basis. To trace the evolutionary history of a candidate gene, a robust phylogenomic framework is indispensable.

**Results:** In this work, we generated a near chromosome-scale genome assembly of the nematode *Rhabditoides inermis* which had previously been proposed as the sister group of the family Diplogastridae to which *P. pacificus* belongs. The genome was assembled from a complex microbial community that consists of multiple bacteria and a fungus of the genus *Vanrija*. The *R. inermis* genome has five chromosomes that likely arose from recent fusions of different Nigon elements. Phylogenomic analysis grouped *R. inermis* within a clade including *C. elegans*, *Mesorhabditis belari* and other rhabditids and thus, did not support a sister group relationship between *R. inermis* and the family Diplogastridae. Comparative genomic analyses identified abundant lineage-specific orthogroups which reveal evidence for parallel expansions of environmentally responsive gene families.

**Conclusions:** Our work demonstrates the value of the *R. inermis* genome as a resource for future phylogenomic analysis and for studying gene family evolution.

## Background

Nematodes are the most abundant animals on Earth and have colonized virtually all ecosystems including multiple animal and plant hosts [1–3]. In addition, with *Caenorhabditis elegans* being a nematode, a vast portion of our biological knowledge originates from basic nematode research. In order to investigate the extent in which biological processes known from *C. elegans* are also conserved in other animals, the free-living nematode *Pristionchus pacificus* had been introduced as a satellite model organism for comparative biology [4]. Over the past years, *P. pacificus* became a more independent model system as new research directions emerged that are unrelated to previous *C. elegans* research. This research focused on an evolutionary novelty of the family Diplogastridae to which *P. pacificus* belongs. Specifically, unlike *C. elegans* and many other nematodes that live in soil habitats and feed on bacteria, *P. pacificus* can form teeth-like denticles in its mouth that allow killing and predation [4]. This predatory form is one of two alternative mouth forms, an example of developmental (phenotypic) plasticity without any intermediate morphologies [5]. The *stenostomatous* (St) morph forms a single tooth with a narrow stoma resulting in strict bacterial feeding. In contrast, the *eurystomatous* (Eu) form has two teeth with a wider stoma, which, in addition to feeding on bacteria, can also kill other nematodes. Research of developmental plasticity and predatory feeding behaviors have established *P. pacificus* as a completely independent model system [6, 7]. The mouth-form dimorphism has become a major model to investigate the molecular mechanisms regulating developmental plasticity, resulting in the identification of a complex gene regulatory network [6, 8–11]. Comparative studies revealed extensive morphological diversification of these feeding structures, highlighting the evolutionary and ecological significance of developmental plasticity. Specifically, more than 50 species of 29 genera of the Diplogastridae family were investigated by morphometric analysis providing multiple examples of structural diversification and evidence for the loss of the dimorphism in individual lineages [12]. Together, these studies have provided a powerful model for investigating plasticity, its molecular causes and organismal significance [13, 14].

The family Diplogastridae to which *P. pacificus* belongs, is a monophyletic taxon that has been subject to intense investigations in the last decade (reviewed in [15–17]). This included the isolation and characterization of new material resulting in the description of six new diplogastrid genera since 2011 [18–23]. These findings suggest that the current phylogenetic and taxonomic knowledge of Diplogastridae is likely still incomplete. In addition, the sister group of the Diplogastridae is unknown and multiple different suggestions have been made over the last two decades. For example, the family Bunonematidae was suggested as a sister group based on morphological comparisons [24] and small subunit (SSU) ribosomal RNA sequencing [25]. In contrast, other rDNA analyses suggested that *Rhabditoides inermis* represents the sister taxon [26–28]. However, a complementary phylogenetic study grouped *R. inermis* and *C. elegans* in a clade of species that are all members of the family Rhabditidae [12]. Recent phylogenomic analysis of the phylum Nematoda did not include *R. inermis* because no genome draft is available [29]. To fill the existing knowledge gap, we here performed genome sequencing of *R. inermis* by PacBio sequencing and Hi-C scaffolding. Using the newly available *R. inermis* genome we performed phylogenomic analysis, which did not provide supporting evidence for a sister group relationship between Diplogastridae and *R. inermis*. Instead, phylogenomic analysis supports a sister group relationship between the Diplogastridae family with a clade of Rhabditidae that includes the genera *Oscheius*, *Caenorhabditis*, *Mesorhabditis*, *Rhabditoides* and *Haemonchus*.

## Results

### *R. inermis* can be repeatedly found in association with the burying beetle *N. vespilloides* in southern Germany

Multiple studies found *R. inermis* to be associated with the burying beetle *Nicrophorus vespilloides* [30, 31]. Burying beetles have a complex biology, e.g. they conceal small vertebrate carcasses underground and prepare these carcasses for consumption by their own offspring [32]. In 2020, we obtained *Nicrophorus vespilloides* beetles from three localities in southern Germany (Bayreuth, Freiburg and Tübingen). We isolated nematodes using standard procedures [33] and obtained more than 12 independent lines that were morphologically similar to the available reference strain *R. inermis* RS5678. Sequencing of the 18S rRNA revealed high similarity and mating experiments confirmed that all strains belong to the same species. Thus, *R. inermis* is found in association with *N. vespilloides* in high frequency and all obtained strains belong to the same species. Also, when using related burying beetle species, which we found in lower frequency, we did not obtain *Rhabditoides* material other than *R. inermis*.

All new isolates of *R. inermis* show a number of features that are different from many other Rhabditidae (including *C. elegans*), but are consistent with original comparative analyses [30]. Specifically, *R. inermis* grows extremely rapidly under laboratory conditions with a generation time of 2-3 days (20° C) when grown on *E. coli* OP50. Distinctively, *R. inermis* is viviparous, resulting in cultures with few if any eggs on the agar plates. Together, we document the consistent association of *R. inermis* with the burying beetle *N. vespilloides* in southern Germany. We established the isolate RS5678 from Great Britain as a reference strain and performed standard inbreeding procedures for 10 generations resulting in a strain RS5678B that was used for genome sequencing. We obtained additional whole genome sequences from two of the german isolates RS6171 and RS6371.

### Genome sequencing data of *R. inermis* represents a complex microbial community

In order to sequence the genome of the inbred *R. inermis* strain RS5678B, we generated genomic DNA libraries from mixed-stage cultures growing on agar plates and sequenced them on the Pacific Biosciences Sequel lI platform. This resulted in 35 Gb of HiFi reads with a median read length of 12 kb. This was assembled into a raw assembly of 2228 contigs spanning 419 Mb (N50=0.4Mb). To reduce the degree of allelism resulting from remaining heterozygosity in the *R. inermis* genome, we ran the HaploMerger2 software on the repeat masked raw assembly [34]. This reduced the number of contigs to 782 spanning 253 Mb with an N50 of 0.9 Mb. We further generated Hi-C data and ran the YaHS software to scaffold the *R. inermis* genome [35]. Visualization of the average GC content and Hi-C coverage for the resulting scaffolds identified five large scaffolds (>10Mb) with a GC content of 47% that span 157Mb. However, a subset of scaffolds also showed an unusually high or low GC content and were mostly sequenced less deeply (Fig. 1A). The Hi-C contact map of the 22 largest scaffolds suggested spatial proximity between the five largest scaffolds, but almost no Hi-C signal between the smaller scaffolds (Fig. 1B). Hypothesizing that the smaller scaffolds could possibly represent microbial contamination on the plates, we annotated protein coding genes and performed a homology search against the NCBI nr database summarizing the phylogenetic distribution of the best hits for all predicted proteins of a given scaffold. This confirmed the microbial origin of these scaffolds (Fig. 1C). Interestingly, none of the microbial contaminants represented *E. coli* OP50 suggesting that all *E. coli* bacteria had been consumed by the time DNA was extracted. Additional sequencing data from two independently isolated *R. inermis* strains showed some sequencing coverage for all identified microbes, suggesting a stable association between *R. inermis* and its microbial community (Additional File 1, Fig. S1).

**Fig. 1.**
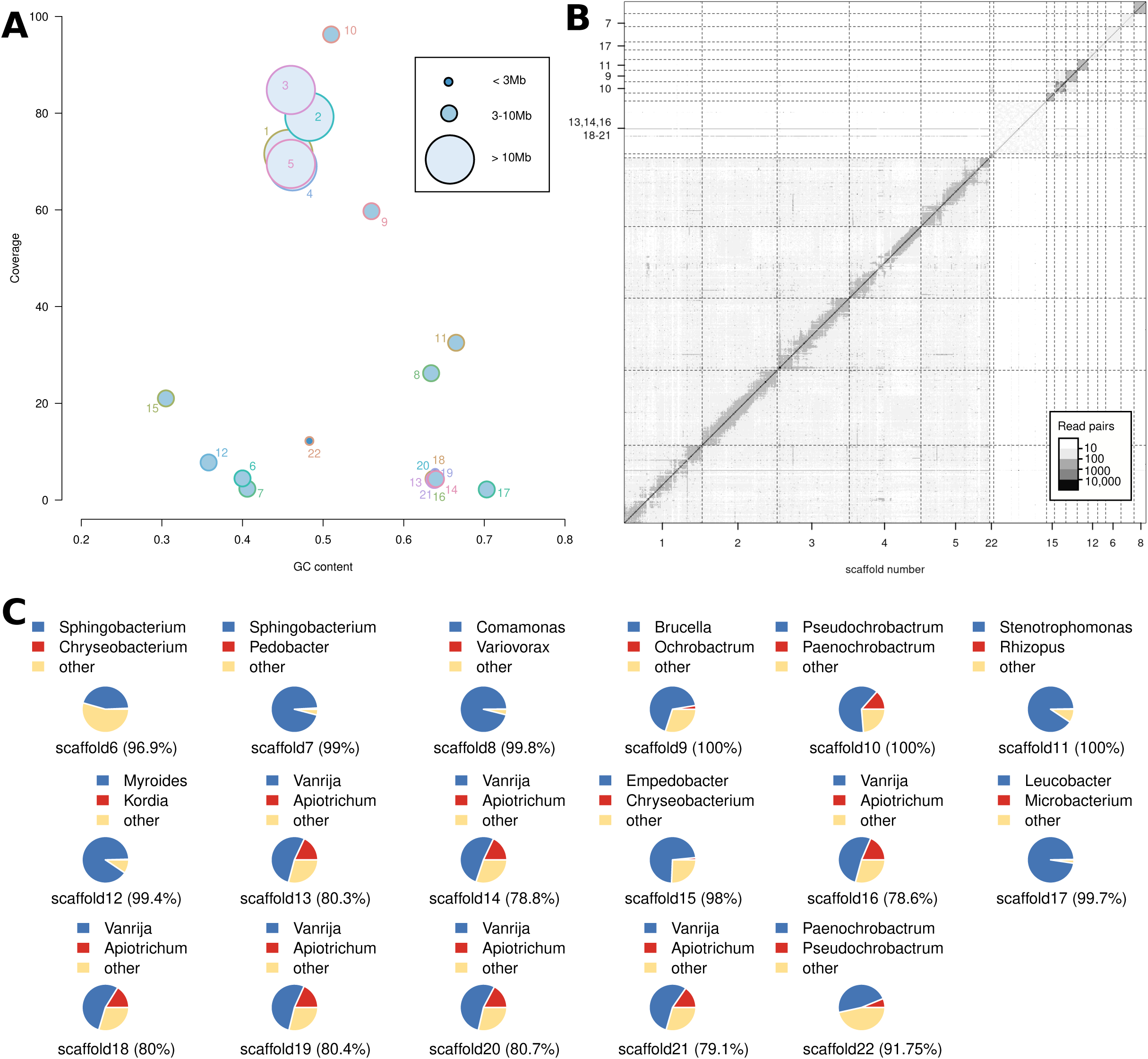
Genome assembly of a complex microbial community. **A** Visualization of GC content and Hi-C coverage indicates that the five largest scaffolds appear to be different from the smaller ones. **B** The Hi-C contact map of the 22 largest scaffolds shows a background level of Hi-C signal across the five largest scaffolds but not between the smaller scaffolds. **C** For scaffolds 6-22 we performed homology searches against the NCBI nr database. The pie charts summarize the taxonomic distribution for the best hits at the genus level.

### A complete fungal genome was assembled from sequencing data of *R. inermis* cultures

While bacterial protein predictions showed a median of more than 90% protein sequence identity with their best hits in the NCBI nr database, multiple scaffolds showed lower identity (∼80%) to fungal sequences of the genus *Vanrija* (Fig. 1C) [36, 37]. This could potentially indicate that our sequencing data represents a novel fungal species or at least that no whole genome data for this species is publicly available. To better characterize our fungal genome (labeled CR7), we first assessed its degree of completeness. The eight scaffolds comprise 22.5Mb of genomic sequence with a GC content of around 65%, which is very comparable to genomic data of other *Vanrija* genomes (Fig. 2). Furthermore, benchmarking of universal single copy orthologs (BUSCO) identified 94.8% of the fungal orthologs (odb9) in our genome assembly [38, 39]. To get more insights into the phylogenetic relationships between the new assembly and publicly available *Vanrija* genomes, we first generated evidence-based gene annotations based on RNA-seq data from *R. inermis* nematode cultures and existing protein annotations of *Vanrija humicola* and *Apiotrichum porosum* [37, 40] and then reconstructed a genome-wide phylogeny based on concatenated alignments of orthologous proteins (Fig. 2, see *Methods*). This confirmed that our fungal genome is closely related but distinct from any other *Vanrija* species, for which whole genome data is available. We then extracted large subunit (LSU) rRNA and internal transcribed spacer (ITS) sequences from the new assembly and performed a combined phylogenetic analysis together with publicly available sequences from diverse *Vanrija* species on NCBI (Additional file 1, Fig. S2). This grouped our sequences together with sequences from *V. albida*. Finally, to further support that our genome most likely represents *V. albida,* we downloaded 56 *V. albida* sequences from NCBI and performed a BLASTN search against our assembly. This showed a median of 100% identity strongly supporting that our genome assembly most likely represents an isolate of *V. albida*. To make this genome publicly available, we deposited it at DDBJ/ENA/GenBank under the accession JAZAQS000000000.

**Fig. 2.**
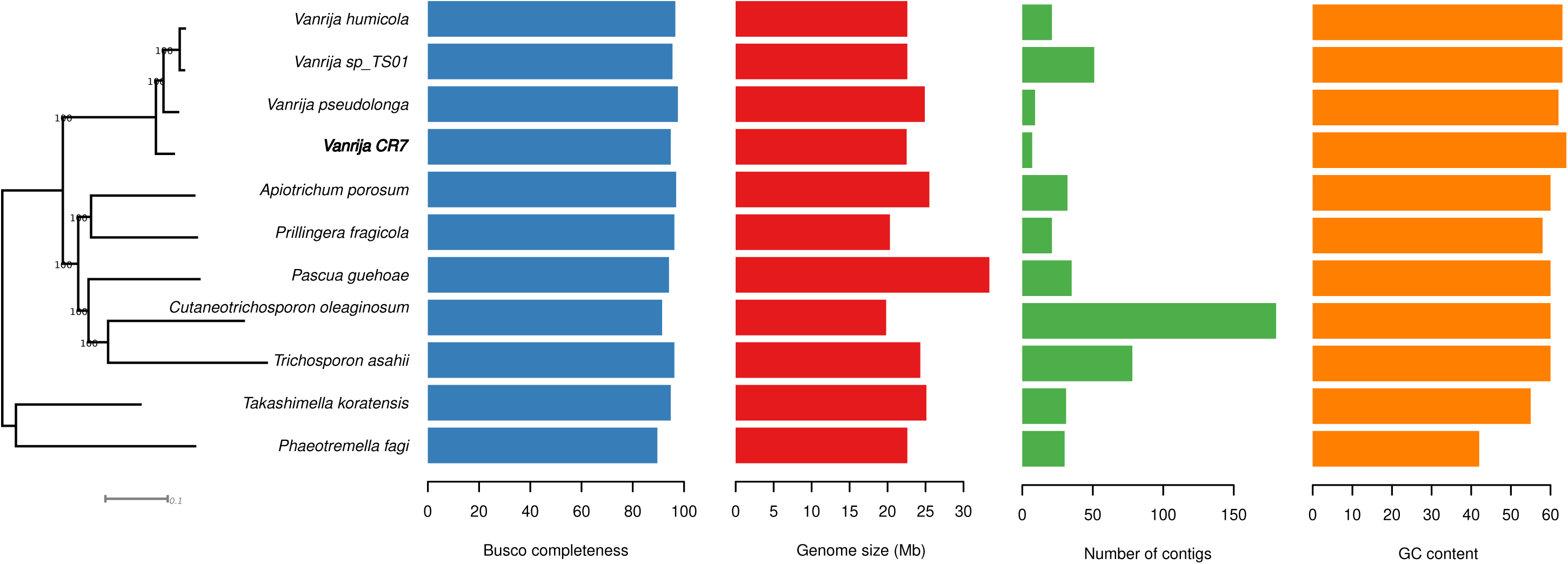
Phylogenomics of fungal genomes from the order Trichosporonales. The phylogeny represents a Maximum-likelihood tree calculated from a concatenated alignment of orthologous proteins. The barplots show the values of various features across the phylogenomic data set.

### *R. inermis* has five chromosomes with near-complete coverage by the five largest scaffolds

To compile a clean version of the *R. inermis* genome, we manually inspected the Hi-C data, GC content, and coverage data for the different scaffolds and performed additional BLASTN searches against the NCBI database. This resulted in a classification of the scaffolds into 213 candidates that are presumably of nematode origin and a subset of scaffolds that represents the microbial community on which the worms were grown. The completeness of the *R. inermis* genome was estimated by BUSCO to be 88.8% (single-copy and duplicates) using the nematode ortholog data set (odb9) [38, 39]. This value is comparable to recently generated nematode genome assemblies from cultures without large-scale microbial contamination [41, 42]. Note that the expected BUSCO completeness value should be around 90%, but will also depend on evolutionary distance from the set of reference species in the ortholog set. The complete *R. inermis* assembly spans 173.4 Mb with an N50 value of 31.4 Mb. The five largest scaffolds comprise 156.7 Mb and thus represent more than 90% of the total assembly. This suggests that *R. inermis* has five chromosomes. To obtain further support for this, we performed a Hoechst staining of the gonads as described previously [42]. This indeed revealed additional evidence for the presence of five chromosomes in *R. inermis* oocytes (Fig. 3A). Thus, genomic data as well as karyotyping analysis support that *R. inermis* has five chromosomes.

**Fig. 3.**
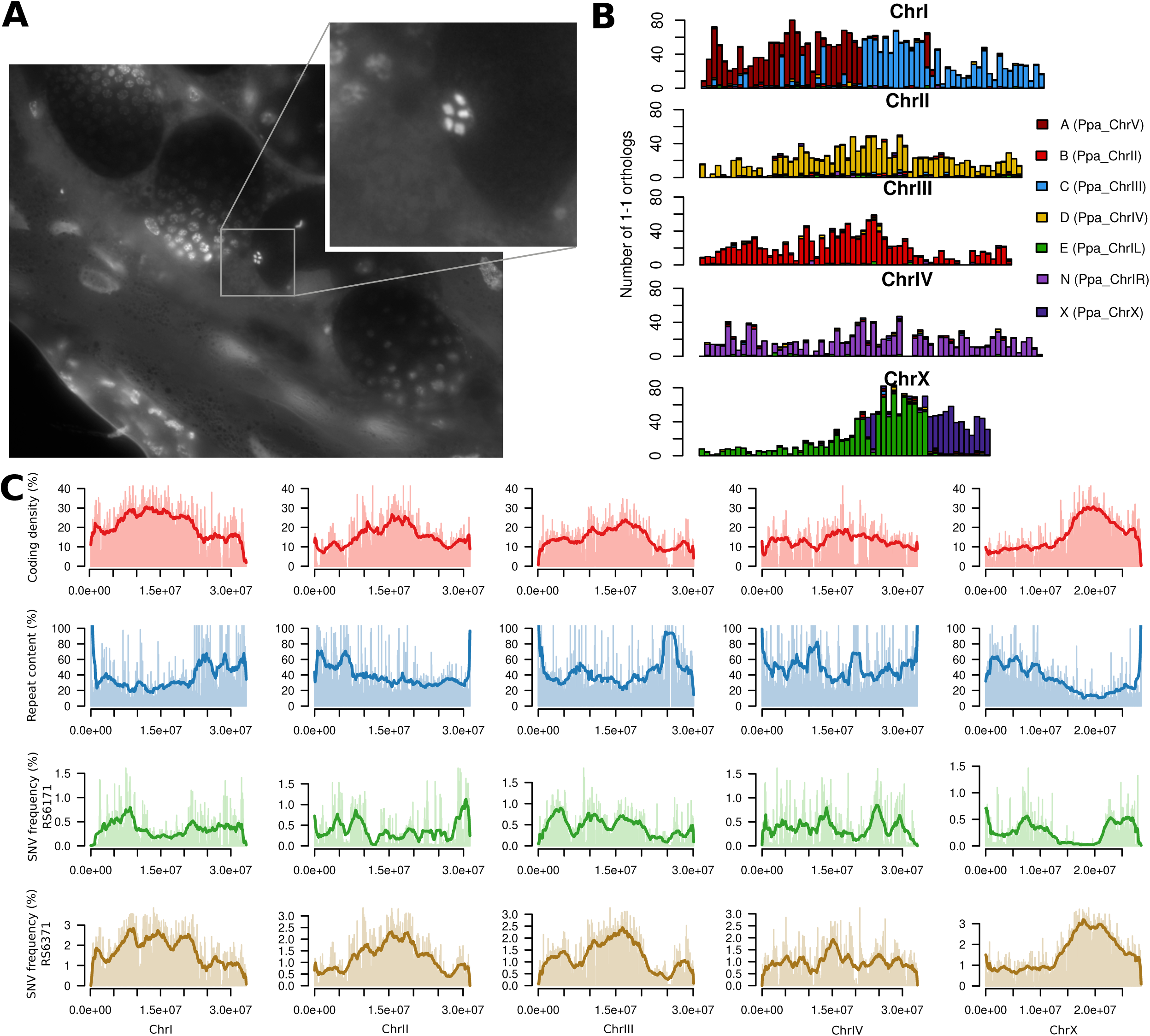
Chromosomal configuration of *R. inermis*. **A** Karyotyping of *R. inermis* worms identified oocytes with five chromosomes. **B** The bars represent 500-kb windows along the five largest scaffolds/chromosomes that show the distribution of 1-1 orthologs color-coded by their chromosomal location in the *P. pacificus* genome which can be used as a proxy to indicate the Nigon elements. **C** The plots show the distribution of coding density, repeat content, and genetic diversity across the *R. inermis* genome. Features were calculated in non-overlapping 100-kb windows and averaged over 20 windows.

### Nigon elements are largely intact in the *R. inermis* genome

The structure of most nematode genomes with chromosome-scale assemblies reflects a specific configuration of seven ancestral linkage blocks that had been called Nigon elements A,B,C,D,E,N and X [43, 44]. These Nigon elements constitute large blocks of macrosynteny between nematode genomes. This means that genes within a Nigon element are frequently reshuffled, whereas rearrangements of genes from different elements are rare. Most nematode chromosomes, specifically in the suborder Rhabditina, correspond to exactly one Nigon element, whereas some other chromosomes are fusions between different Nigon elements. Examples of fused chromosomes are the *Caenorhabditis* chromosome X, *P. pacificus* chromosome I and many others [44–47]. The observation of five chromosomes in *R. inermis* suggests that two fusions of Nigon elements occurred. We generated evidence-based gene annotations based on protein homology and transcriptome data with a BUSCO completeness of 86%. We used these protein annotations to visualize Nigon elements across the *R. inermis* genome, which revealed one fusion of Nigon element A and C resulting in *R. inermis* chromosome I and a second fusion between elements E and X resulting in *R. inermis* chromosome X (Fig. 3B). Note that we labeled chromosome X based on the observation that Nigon element X tends to form the X chromosome [44]. However, all chromosomes have very similar coverage (Fig. 1A), which makes the identification of a good candidate for a sex chromosome difficult. Based on these observations, we would speculate that the sex chromosome of *R. inermis* was formed recently and there is very little differentiation between the X and Y chromosome.

### Analysis of coding density, repeat content and genetic diversity reveals evidence for the presence of chromosome centers

Many nematode chromosomes also display an internal structure with chromosomal centers and arms [45, 48]. As the chromosomes of *C. elegans* and *P. pacificus* are holocentric [49, 50], *i.e.* centromeric function is distributed along the whole length of the chromosomes, chromosome centers are not centromeres. The chromosome centers are rather distinguished by higher gene density and a bias towards older gene classes, lower recombination, less genetic diversity, and lower repeat content as opposed to the chromosome arms [45]. These patterns are thought to be generated by higher levels of background selection in the centers [51]. It has to be noted, that the distinction between arms and centers is not universal across nematode chromosomes and it has been shown that chromosome fusions with associated shifts in the recombinational landscapes can drastically alter intra-chromosomal structures [47]. To test for the presence of center-like regions in the *R. inermis* genome, we calculated the coding density (percentage of protein coding sequence), repeat content and genetic diversity in 100-kb windows (Fig. 3C). This revealed some central peaks of high coding density that coincided with low repeat content on chromosomes I,II,III, and X. With regard to genetic diversity, the pattern becomes less clear. While the more closely related strain RS6171 also shows lower SNP density in central regions of chromosomes I,II, and X, these patterns appear to be reversed in the more divergent strain RS6371 (Fig. 3C). This could potentially be explained by different recombinational landscapes between the strains and could be tested in future by experimental crosses. Nevertheless, the genetic diversity in the more closely related RS6171 together with the analysis of coding density and repeat content point towards the presence of center-like regions on *R. inermis* chromosomes I, II, and X.

### Phylogenomic analysis does not support *R. inermis* as a sister taxon to Diplogastridae

Previous phylogenetic studies suggested a potential sister taxon relationship between *R. inermis* and the family Diplogastridae [26, 27]. To resolve the phylogenetic relationship between *R. inermis* and diplogastrid nematodes, we compiled a phylogenomic data set of annotated proteins and transcriptomes from 40 species [41, 42, 51–69]. We employed a previously established phylogenomic pipeline for reconstructing a species tree from this type of data [70]. This resulted in a concatenated alignment of 1402 single-copy orthologous genes that was taken to compute a maximum-likelihood tree (Fig. 4, see *Methods*). This phylogeny groups *R. inermis* within a clade of rhabditids including *C. elegans* and *M. belari*. Thus, this grouping did not support the position of *R. inermis* as sister group to the diplogastrids. Importantly, the genera *Poikilolaimus* and *Bunonema* are excluded from this clade and appear as next outgroups. In order to explore potential reasons for the misplacement of *R. inermis* in previous phylogenies, we reanalyzed available nematode sequences for the RNApolII and the 18S and 28S ribosomal RNA markers. Of all three markers, only the 18S sequences supported a sister group relationship (Additional file 1, Fig. S3). This suggests that previous phylogenetic analysis is reproducible and that the 18S sequence largely contributed to the previous misplacement of *R. inermis*.

**Fig. 4.**
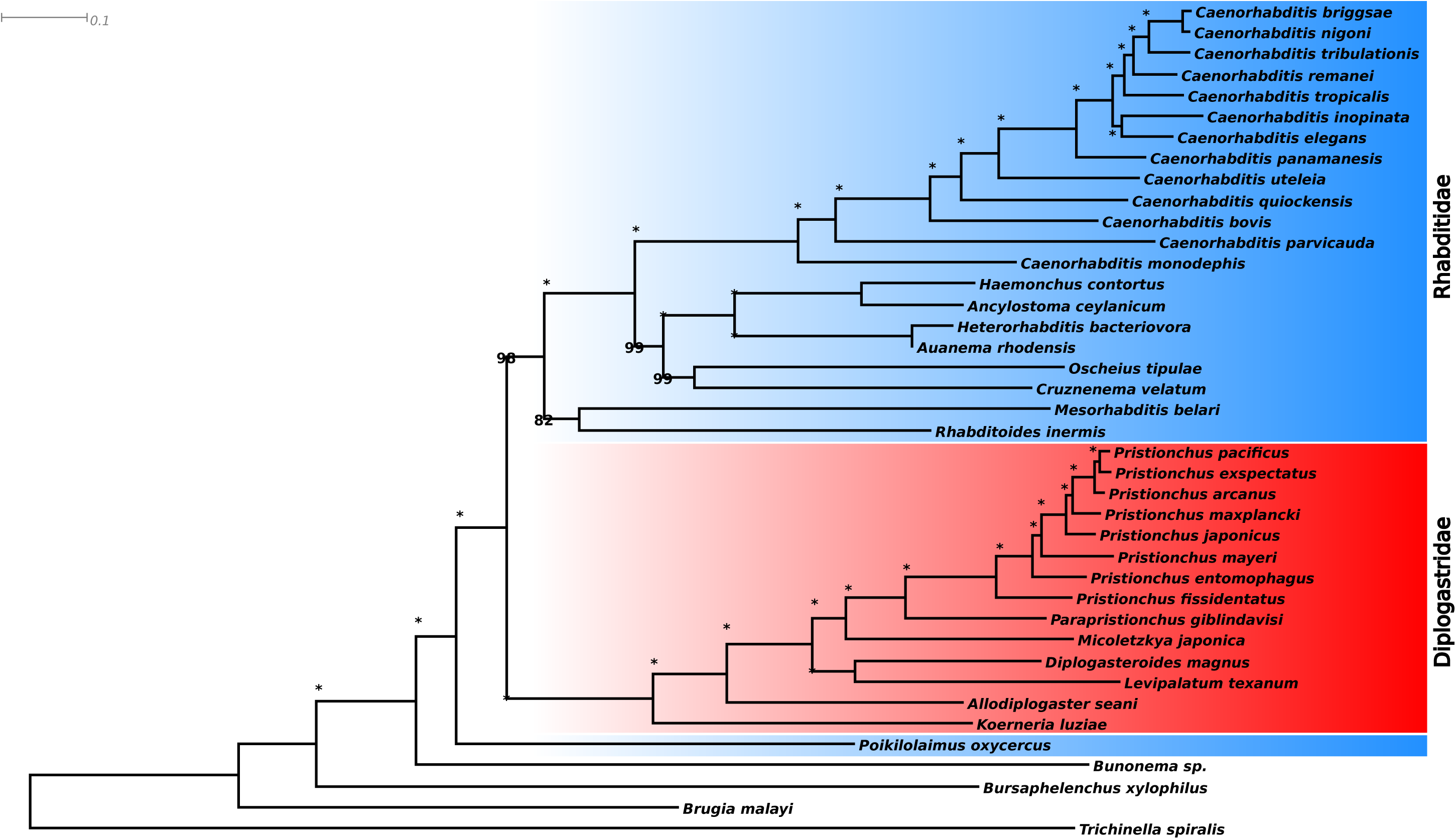
Phylogenomics of nematodes of the suborder Rhabditina. Phylogenomic and transcriptomic data of diplogastrid and rhabditid nematodes were used to reconstruct a maximum-likelihood tree for the suborder Rhabditina. Note that the family Rhabditidae is known to be paraphyletic [25]. The species *B. xylophilus*, *B. malayi,* and *T. spiralis* were included as outgroups. Stars indicate a bootstrap support of 100/100.

### Comparative genomic analysis reveals parallel gene family expansions across multiple nematode lineages

To further characterize the gene set of *R. inermis*, we performed an orthology analysis with *C. elegans*, *P. pacificus*, *M. belari*, *H. contortus*, and *O. tipulae*. The program Orthofinder identified 17,464 orthogroups with a minimum size of two genes. 6135 orthogroups have at least one ortholog in all six species (Fig. 5A). The presence of orthologs in all six species is the most abundant pattern among all orthogroups. Orthogroups reflecting species-specific absence or presence represent the next most frequent patterns (Fig. 5A). This could be expected since none of the species are closely related and the terminal branches in the underlying species tree are relatively long. As the minimum size of an orthogroup is two, species-specific orthogroups can be interpreted as lineage specific duplication events. Thus, the number of 1752 orthogroups that are specific to the *R. inermis* branch translates to 5985 *R. inermis* genes. We tried to further characterize this gene set by screening for overrepresented protein domains. This revealed a drastic expansion of genes with a BTB domain and an associated Kelch domain (Fig. 5B). These domains are typically found in adapter proteins regulating protein degradation. In *P. pacificus*, BTB domain containing proteins were found to be enriched in the germline [71] and multiple studies reported a loss of BTB genes following the transition to hermaphrodites [42, 62, 70, 72]. The expansion of this gene family in the *R. inermis* lineage apparently occurred in parallel to the expansions in the lineages leading to *C. elegans* and *Panagrellus redivivus* [73, 74]. This could imply that the expansion of the BTB gene family may be a general pattern in nematodes. To test if even more lineages share the gene expansions of BTBs and other gene families, we repeated this analysis with all other sets of species-specific orthogroups. This revealed 22 protein domains that show significant enrichments in at least two lineages (Bonferroni corrected P-value < 0.05, Fig. 5C). While no gene family is significantly enriched in all six species, G-protein coupled receptors (GPCRS) show up in five lineages. Furthermore BTB, F-box, and C-type lectins show significant enrichments in four lineages. This suggests that multiple gene families underwent parallel expansions in different nematode lineages.

**Fig. 5.**
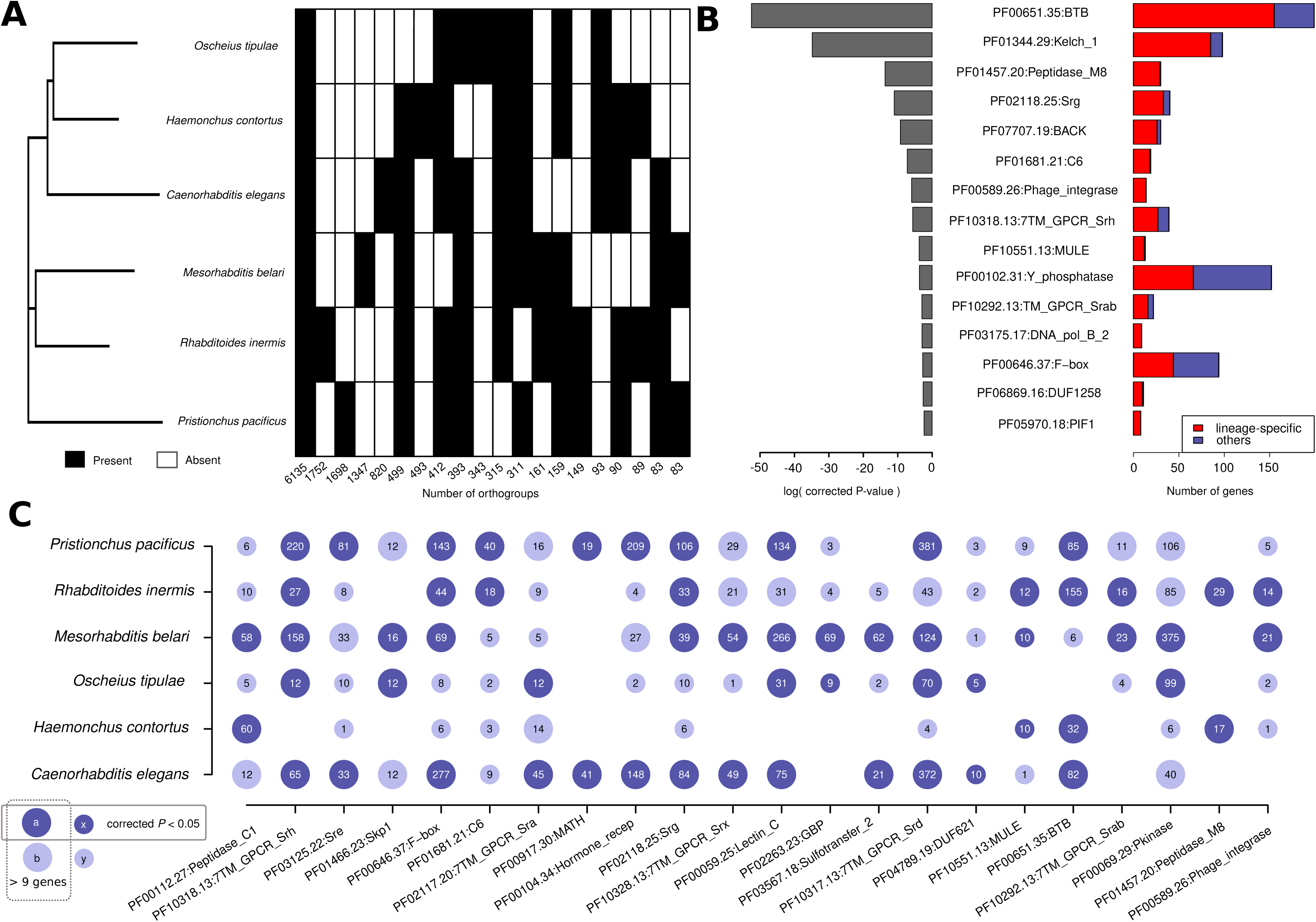
Parallel gene family expansions across multiple nematode lineages. **A** Orthogroups were computed by the OrthoFinder pipeline and classes of orthogroups were defined based on the absence/presence patterns and were sorted by abundance. Orthology classes are visualized as a heatmap next to the phylogeny. With exception of the first class of 6135 orthogroups that are shared between all species, the most abundant patterns mostly reflect lineage-specific patterns such as species-specific orthogroups or orthogroup that are only missing in a single data set. **B** The barplots indicate the significance of enrichment and the total number of *R. inermis* genes with protein domains. **C** We repeated the protein domain enrichment analysis for all species-specific orthogroups. The bubble plot visualizes the significance and the number of genes in species-specific orthogroups per domain and species.

## Discussion

Free-living nematodes like *C. elegans* and *P. pacificus* are not only powerful model systems for genetic studies, they are also an excellent subject for evolutionary genomic studies as their species richness allows us to test various hypotheses across different clades and multiple time-scales [51, 72]. Two major factors are essential for such studies. First, deep taxon sampling is needed to facilitate comparisons at high phylogenetic resolution. Over the last two decades, continued sampling efforts have increased the number of known species in the genera *Caenorhabditis* and *Pristionchus* from 10 and 18, respectively to over 50 for both genera [59, 75–77]. Second, we need a robust phylogenetic framework to correctly infer the evolutionary history of genomic features. Abundant genomic and transcriptomic resources have helped to resolve various ambiguities about the phylogeny of the nematode phylum [29, 78, 79]. In this context, the sequencing of the *R. inermis* genome was needed to clarify the question about the sister group to the family Diplogastridae. Our phylogenomic analysis does not support *R. inermis* as sister to Diplogastridae and re-analysis of previous data suggests that the misplacement of *R. inermis* is largely due to phylogenetic signals coming from the 18S rRNA sequence. Although the phylogenetic position of *R. inermis* makes its genome not particularly interesting for studies focusing on *P. pacificus*, we demonstrate its value for detecting parallel gene family expansions across multiple lineages. It should be noted that additional analysis using more deeply sampled taxa is needed to pinpoint the exact timing of the expansions. Thus, a more highly resolved study in diplogastrid nematodes dated the origin of many BTB orthologs to the ancestor of the *Pristionchus* genus [80]. Given that GPCRs are expressed in chemosensory neurons [81], C-type lectins were shown to carry out important functions in the immune response [82], and also F-box and BTB proteins are also thought be involved in the response to pathogens [73], all identified gene families seem to be involved in the response to environmental stimuli. Thus, it is tempting to speculate that the dynamic evolution of these gene families reflects adaptation to new environments. *R. inermis* has a tight association with the burying beetle *N. vespilloides.* Note that *Rhabditoides* is the only nematode genus known to be reliably associated with burying beetles. Our genome data suggests that this ecological interaction also involves a complex microbial community containing bacteria and fungi. Currently, we do not know which of the microbes forms a component of the nematode’s gut microbiome and whether there are any synergistic effects between different species [83]. Thus, the *R. inermis* associated microbial community might be an interesting system to investigate complex interplay between nematodes, bacteria, and fungi.

## Conclusions

The main motivation of this study was the identification of the potential sister group of the nematode family Diplogastridae that has long been debated. Previous genomic studies remained inconclusive because not all known taxa had available genome drafts. With this study on *Rhabditoides* one knowledge gap is filled. This work, and the new phylogeny provided in here, supports a sister group relationship between the monophyletic Diplogastridae and a group of genera of Rhabditidae including *C. elegans* and *R. inermis*. In contrast to previous phylogenetic analysis, no single genus or species shows a sister group relationship with Diplogastridae and instead, a sub-clade of rhabditids with substantial morphological and ecological diversification appears as a sister taxon. While the future identification and characterization of additional, currently unknown taxa, might change this phylogeny, the new pattern is consistent with the massive radiation and ecological diversification of two major nematode lineages in soil and associated habitats.

## Methods

### Nematode culturing and preparation of sequencing libraries

Nematodes of *R. inermis* were grown on standard nematode growth medium as for *C. elegans* and *P. pacificus* [84]. Worms were washed off of 50 fully grown plates using M9 buffer, filtered by using a 5µm filter in order to remove remaining bacteria and then gently pelleted by centrifugation (1,300 g, 1min). Pelleted worms were frozen in liquid nitrogen and ground to a fine powder using a mortar and pestle. The powder was directly transferred into the lysis buffer used in combination with Qiagen genomic tip columns (500/G) (Qiagen, Hamburg, Germany). The protocol was performed following the manufacturer’s instructions. All steps involving sample vortexing were replaced by sample inverting to limit unwanted DNA shearing. DNA quality and quantity were determined with a NanoDrop ND 1000 spectrometer (PeqLab, Erlangen, Germany), a Qubit 2.0 Fluorometer (ThermoFisher Scientific, Waltham, USA), and by a Femto pulse system (Agilent, CA, USA). A total of 10 µg genomic DNA was sheared to a target fragment size of 19 kb using a Megaruptor 2 device (Diagenode, Denville, USA). In order to remove small fragments (<8kb), the BluePippin size-selection system was used according to the manufacturer’s protocol (P/N 100-286-000-07, Pacific Biosciences, USA). The final library was sequenced on a SMRT cell of a Pacific Biosciences Sequel II instrument following the Magbead loading protocol. RNA-seq data of mixed-stage cultures was generated as described previously. For Hi-C analysis, *R. inermis* nematodes were grown on standard nematode growth medium and washed off with M9 Buffer. 50µl frozen worm pellet was ground with a motor in an Eppendorf tube and used for Hi-C Library preparation following the manufacturer’s instructions for small animals (Arima Genomics, California, USA). RNA-seq libraries were prepared from mixed-stage cultures as described previously [70].

### Genome assembly

Raw PacBio reads were converted into Hifi reads by the ccs program (version 6.4.0) with –min-rq 0.99 option and then demultiplexed by the lima program (version 2.6.0). The Hifi reads were then assembled by the Canu program (version v1.4) with options: -pacbio-corrected -genomeSize=400m [85]. Haplotypes were merged by the program Haplomerger2 (version 20180603) after softmasking repeats in the draft assembly with the help of the Red software (version 05/22/2015) [86, 87]. Hi-C data was aligned against the haplomerged assembly by BWA mem (version 0.7.17-r1188) and scaffolded using the yahs tool (version 1.2a.2) [35, 88]. For metagenomic classification, we annotated scaffolds with Prokka (version 1.14.6) and performed DIAMOND (version 2.1.4.158) searches against the NCBI NR database (downloaded August 2023) [89, 90]. The taxonomic distributions of the best hits were summarized using the taxonomizer library in R. Based on manual inspection of Hi-C data, GC content, coverage data, and additional BLASTN searches against the NCBI database, we classified the scaffolds into 213 candidates that presumably are of nematode origin and a subset of scaffolds that represents the microbial community on which the worms were grown. At this step, we relabeled the five largest scaffolds as chromosomes (ChrI-ChrX) and manually extracted some regions from scaffolds 1 and 4 that showed some Hi-C signal and merged them as a separate scaffold (ChrUn). Karyotyping analysis was performed as described previously [42]. In order to visualize the repeat content across the *R. inermis* genome, the software Red was run to detect repetitive sequences in the final assembly.

### Gene annotation

A *de novo* transcriptome assembly was assembled from mixed-stage RNA-seq data using trinity (version 2.2.0) with the normalize_reads option [91]. This transcriptome assembly and the current *P. pacificus* gene annotations (version El Paco gene annotations 3 [92]) were used to generate a set of evidence-based gene annotations using the PPCAC pipeline [93]. In short, transcribed open reading frames and proteins were mapped against the *R. inermis* genome by the exonerate protein2genome tool (version 2.2.0) with options: --bestn 2 --dnawordlen 20 --maxintron 20000 [94]. Subsequently, per locus (100-bp window) and strand, the gene model with the longest ORF was chosen as representative gene model. We then applied a filtering step to remove spurious annotations that likely result from falsely called ORFs in our transcriptome data. Such spurious annotations will result in species-specific orphan genes (SSOGs) without protein homology in other genomes that are located in the antisense direction of conserved genes, i.e. genes with homologs in other species [80, 92]. In short, *R. inermis* SSOGs were defined by blastp searches against the protein sets of *C. elegans*, *O. tipulae*, *P. pacificus*, *H. contortus*, and *M. belari* (e-value < 0.001) and SSOGs overlapping conserved genes on the antisense strand were removed from the annotation. This resulted in 23,442 gene models with a BUSCO (version 3) completeness score of 85.9% (complete + duplicated, against the odb9 nematode data set) [38, 39]. Coding densities, i.e. the percentage of protein-coding sequences within 100-kb windows were extracted from the resulting annotation files. For coloring the Nigon elements, best reciprocal BLAST hits were computed by the get_BRH.pl of the PPCAC pipeline using the pairwise BLAST output files between *R. inermis* and *P. pacificus* (e-value <0.001).

### Population genomic analysis

Raw sequencing data for the *R. inermis* strains RS6171 and RS6371 were aligned against the *R. inermis* genome by the BWA mem alignment program (version 0.7.17-r1188). Single nucleotide variants (SNVs) calls were generated by mpileup and vcfutils.pl varFilter (options: -w0 -D1000) commands of the samtools software suite (version 0.1.8). Only variants with a quality score of at least 20 were taken for visualization of genetic diversity across the *R. inermis* genome.

### Phylogenomic species tree

For most species, we obtained protein-coding gene predictions from WormBase ParaSite (Table S1) [95]. For genes, with multiple annotated isoforms, we used the longest protein as a representative sequence. For other species, we generate protein predictions from available transcriptome data. In such cases, either transcriptome assemblies were downloaded from the European Nucleotide Archive or were generated by the Trinity assembler (version 2.2.0 with –normalize_reads option). Subsequently, the longest partial or complete open reading frame was called per sequence and potential redundant sequences resulting from alternative splicing were removed by running the cd-hit clustering program (version 4.3 with default 90% sequence identity threshold). For the reconstruction of a species tree, we followed a similar procedure as described previously [70]. In short, orthologous groups were calculated by the orthagogue [96]. Orthogroups with at most one sequence per species were taken to generate multiple sequence alignments using the MUSCLE program (version 3.8.1551). The concatenated alignments were taken as input to the RAxML software (version 8.2.12) which reconstructed a maximum-likelihood tree (options: -m PROTGAMMAILG -f a ) [97].

### Orthology and protein domain enrichment

We ran the OrthoFinder pipeline (version 2.5.2) [98] on the *R. inermis*, *M. belari*, *P. pacificus*, *C. elegans*, *O. tipulae*, and *H. contortus* data sets (Additional File 1, Table S1) and extracted sets of orthogroups with specific absence and presence patterns from the Orthogroups output folder. To test for overrepresented protein domains among species-specific orthogroups, we scanned for protein domains with the hmmsearch program (Version 3.3 with -R 0.001 option) using the Pfam-A database (version 3.3) [99, 100]. Tests for overrepresentation were performed by Fisher’s exact-test with Bonferroni correction for multiple testing.

### Gene tree analysis

For each tested marker (ITS and LSU from fungi and RNApolII, 18S, 28S from nematodes ), we downloaded corresponding sequences for representative species from NCBI and generated a multiple sequence alignment with MUSCLE (version 3.8.1551) [101]. Phylogenetic trees were calculated using the R phangorn package [102]. Specifically, the modelTest function was run on each individual data set and a final maximum likelihood tree was calculated using bootstrap.pml function selecting the best model based on the Bayes information criterion.

## Abbreviations

ITS: internal transcribed spacer
LSU: large subunit ribosomal RNA
SNV: Single nucleotide variant
SSOGs: Species-specific orphan gene
SSU: small subunit ribosomal RNA

## Declarations

### Ethics approval and consent to participate

Not applicable

### Consent for publication

Not applicable

### Availability of data and materials

The genome assembly and sequencing reads have been uploaded to the European Nucleotide archive under the study accession PRJEB71643. The fungal genome of the genus *Vanrija* has been deposited at DDBJ/ENA/GenBank under the accession JAZAQS000000000.

### Authors’ contributions

Conceptualization, R.J.S.; Investigation, C.R., W.R., M.A., S.W., M.H. and R.J.S.; Data curation, C. R.; Visualization, M.A., S.W. and C. R.; Writing original draft, C.R.; Writing – review & editing, C.R. and R.J.S; Project administration, C.R. and R.J.S.; Supervision, C.R and R.J.S.

## Acknowledgements

We would like to thank the whole Sommer lab for helpful discussions.

## Supplementary Information Additional file 1

**Fig. S1:** Coverage of microbial scaffolds in different *R. inermis* isolates.

**Fig. S2.** Phylogenetic relationships between different *Vanrija* species.

**Fig. S3.** Phylogenetic relationships between different nematodes.

**Table S1** Nematode genomic and transcriptomic data sets.

