## Supplemental figures and tables for "The genome assembly of *Rhabditoides inermis* from a complex microbial community reveals further evidence for parallel gene family expansions across multiple nematodes"

[2] Current address: Institute of Molecular Biotechnology of the Austrian Academy of Sciences (IMBA), Vienna BioCenter (VBC), 1030 Vienna, Austria.

[*] Corresponding authors:

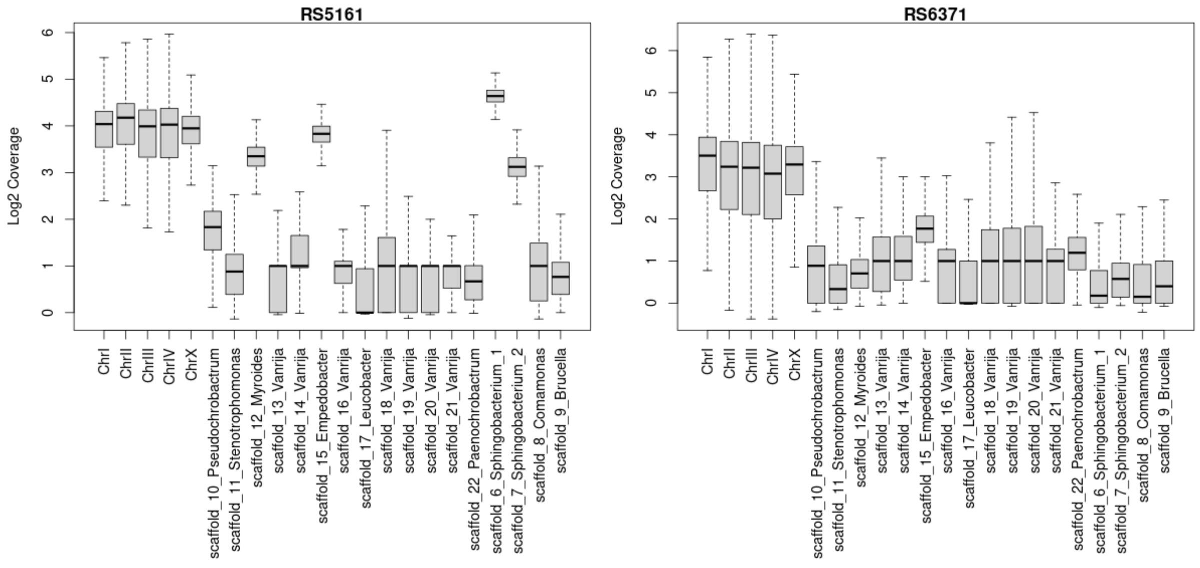


**Fig. S1. Coverage of microbial scaffolds in different *R. inermis* isolates.** Coverage analysis reveals evidence for the presence of all microbes in sequencing data of two independent isolates of *R. inermis*. The y-axis indicates the average Log2 coverage in 1-kb windows for the largest scaffolds/chromosomes.

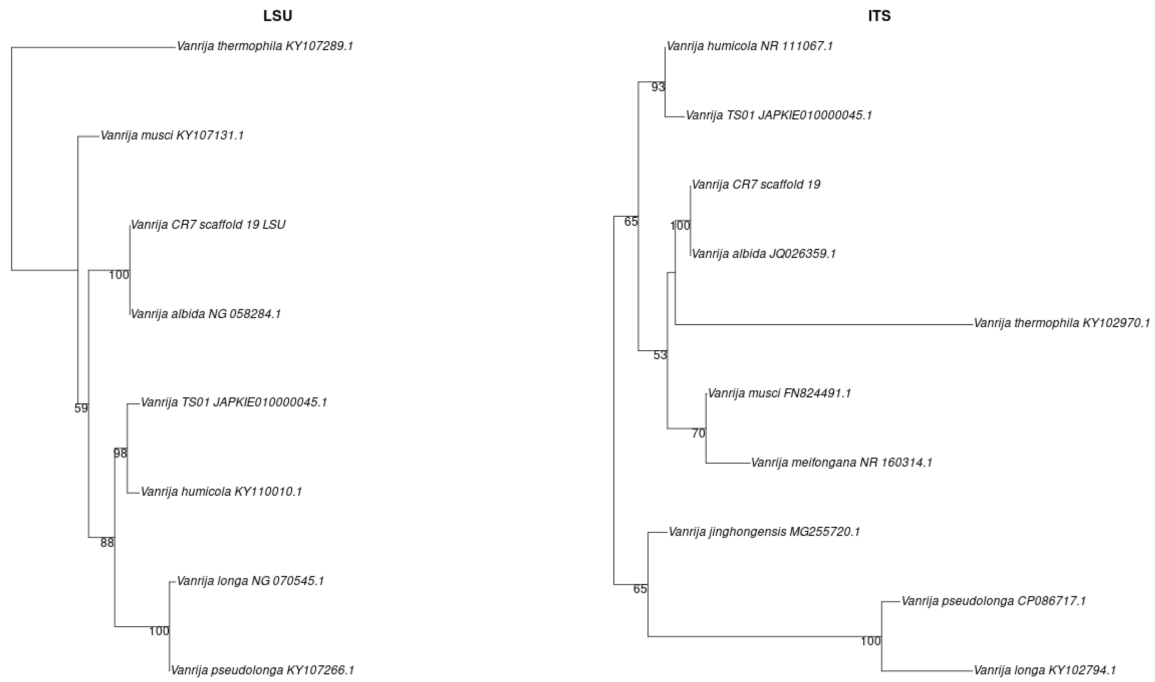


**Fig. S2. Phylogenetic relationships between different *Vanrija* species.** LSU and ITS sequences from different *Vanrija* species were downloaded from NCBI Genbank, aligned with MUSCLE, and maximum likelihood trees were calculated by the phangorn R package. For both markers, sequences from our genome assembly (CR7) group together with sequences from *V. albida*.

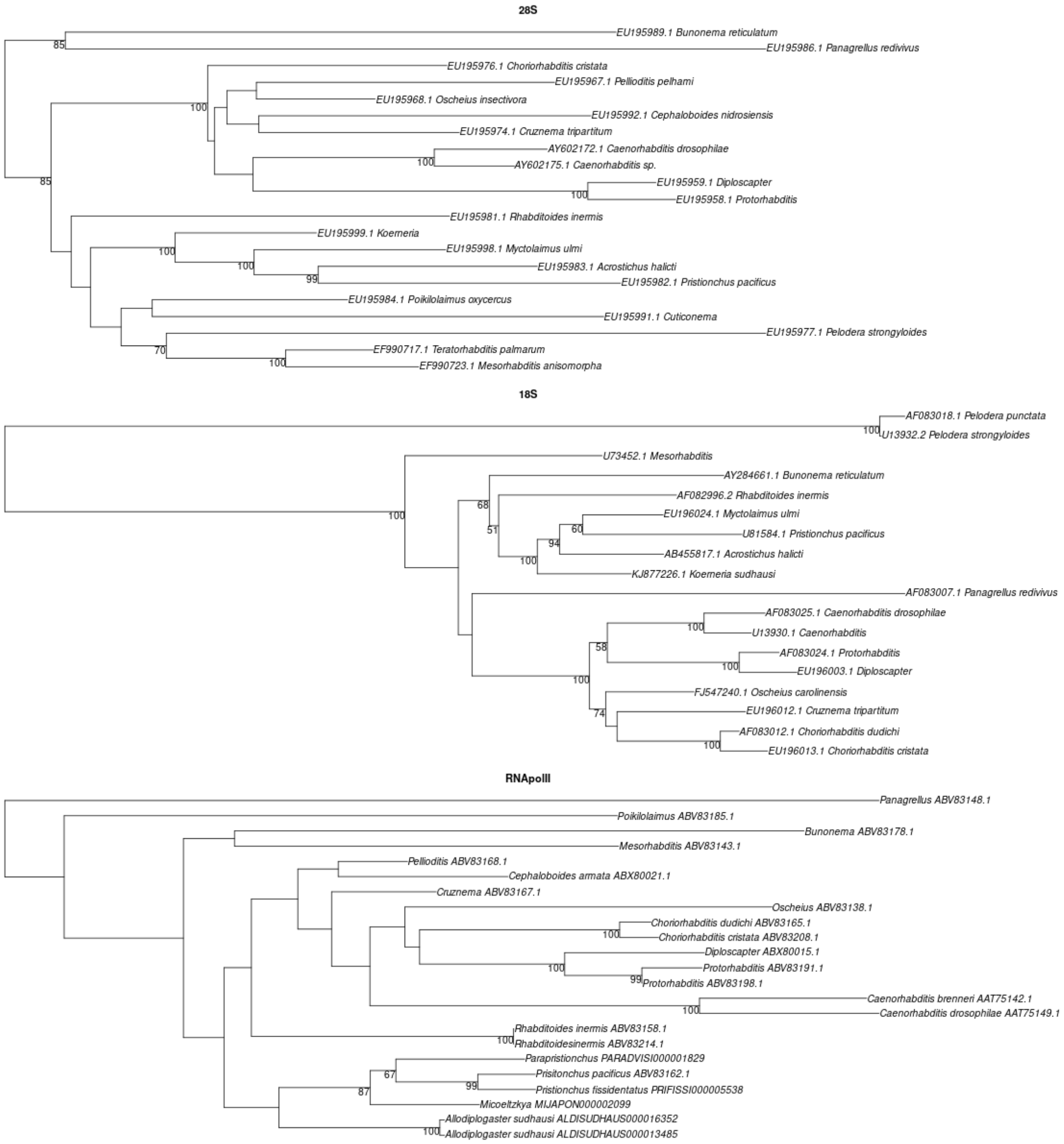


**Fig. S3. Phylogenetic relationships between different nematodes.** Phylogenetic trees were reconstructed for RNApolII, 18S and 28S ribosomal RNAs from different nematodes. Only the 18S data suggests a sister group relationship between R. inermis and the family Diplogastridae.

### Table S1 - Nematode genomic and transcriptomic data sets

| Species | Source accession / version | Type | Reference |
|---|---|---|---|
| *Allodiplogaster seani* | European Nucleotide Archive: HCAZ01000000 | RNA | Wighard et al. 2022 |

| | | | |
|---|---|---|---|
| *Ancylostoma ceylanicum* | WormBase ParaSite WBPS18 PRJNA231479 | Protein | Schwarz et al. 2015 |
| *Auanema rhodensis* | European Nucleotide Archive: ERR3150287 | RNA | Tandonnet et al. 2019 |
| *Bunoema sp.* | European Nucleotide Archive: GITZ01000000 | RNA | Casasa et al. 2021 |
| *Brugia malayi* | WormBase ParaSite WBPS14 PRJNA10729 | Protein | Ghedin et al. 2007 |
| *Bursaphelenchelus xlyophilus* | WormBase ParaSite WBPS14 PRJEA64437 | Protein | Kikuchi et al. 2011 |
| *Caenorhabditis bovis* | WormBase ParaSite WBPS18 PRJEB34497 | Protein | Stevens et al. 2019 |
| *Caenorhabditis briggsae* | WormBase ParaSite WBPS14 PRJNA10731 | Protein | - |
| *Caenorhabditis elegans* | WormBase ParaSite WBPS14 PRJNA13758 | Protein | - |
| *Caenorhabditis inopinata* | WormBase ParaSite WBPS18 PRJDB5687 | Protein | Kanzaki et al. 2018 |
| *Caenorhabditis monodelphis* | caenorhabditis.org | Protein | Slos et al. 2017 |
| *Caenorhabditis nigoni* | WormBase ParaSite WBPS18 PRJNA384657 | Protein | Yin et al. 2018 |
| *Caenorhabditis panamensis* | WormBase ParaSite WBPS18 PRJEB28259 | Protein | Stevens et al. 2019 |
| *Caenorhabditis parvicauda* | WormBase ParaSite WBPS18 PRJEB12595 | Protein | Stevens et al. 2019 |
| *Caenorhabditis quiockensis* | WormBase ParaSite WBPS18 PRJEB11354 | Protein | Stevens et al. 2019 |
| *Caenorhabditis remanei* | WormBase ParaSite WBPS18 PRJNA248911 | Protein | Fierst et al. 2015 |
| *Caenorhabditis tribulationis* | WormBase ParaSite WBPS18 PRJEB12608 | Protein | Stevens et al. 2019 |
| *Caenorhabditis tropicalis* | WormBase ParaSite WBPS18 PRJNA53597 | Protein | - |
| *Caenorhabditis uteleia* | WormBase ParaSite WBPS18 PRJEB12600 | Protein | Stevens et al. 2019 |
| *Cruznema velatum* | European Nucleotide Archive: SRR23934717 | RNA | Guo et al. 2023 |
| *Diplogasteroides magnus* | European Nucleotide Archive: GITX01000000 | RNA | Casasa et al. 2021 |

| | | | |
|---|---|---|---|
| *Haemonchus contortus* | WormBase ParaSite WBPS14 PRJEB506 | Protein | Doyle et al. 2020 |
| *Heterorhabditis bacteriovora* | European Nucleotide Archive: SRR6294669 | RNA | Vadnal et al. 2018 |
| *Koerneria luziae* | European Nucleotide Archive: GIUA01000000 | RNA | Casasa et al. 2021 |
| *Levipalatum texanum* | European Nucleotide Archive: GITY01000000 | RNA | Casasa et al. 2021 |
| *Micoletzkya japonica* | pristionchus.org | Protein | Prabh et al. 2018 |
| *Oscheius tipulae* | WormBase ParaSite WBPS14 PRJEB15512 | Protein | Besnard et al. 2017 |
| *Poikilolaimus oxycercus* | European Nucleotide Archive: SRR6049087 | Protein | Beltran et al. 2018 |
| *Pristionchus arcanus* | pristionchus.org PPCAC (version 1) | Protein | Prabh et al. 2018 |
| *Pristionchus entomophagus* | pristionchus.org PPCAC (version 1) | Protein | Prabh et al. 2018 |
| *Pristionchus exspectatus* | pristionchus.org Yoshida et al. | Protein | Yoshida et al. 2023 |
| *Pristionchus fissidentatus* | pristionchus.org PPCAC (version 1) | Protein | Prabh et al. 2018 |
| *Pristionchus japonicus* | pristionchus.org PPCAC (version 1) | Protein | Prabh et al. 2018 |
| *Pristionchus mayeri* | pristionchus.org PPCAC (version 1) | Protein | Prabh et al. 2018 |
| *Pristionchus maxplancki* | pristionchus.org PPCAC (version 1) | Protein | Prabh et al. 2018 |
| *Pristionchus pacificus* | pristionchus.org, El Paco gene annotation 3 | Protein | Athanasouli et al. 2020 |
| *Parapristionchus giblindavisi* | pristionchus.org (version 2, 2022) | Protein | Röseler et al. 2022 |
| *Rhabditoides inermis* | this study | Protein | this study |
| *Trichinella spiralis* | WormBase ParaSite WBPS14 PRJNA12603 | Protein | Mitreva et al. 2011 |
